# Fluorescence correlation spectroscopy and photon counting histograms in small domains. Part I: General theory

**DOI:** 10.1101/847129

**Authors:** Y. Jiang, B. Xu, A. Melnykov, G. M. Genin, E. L. Elson

## Abstract

Analysis of fluctuations arising as fluorescent particles pass through a focused laser beam has enabled quantitative characterization of molecular kinetic processes. The mathematical frameworks of both fluorescence correlation spectroscopy (FCS) and photon counting histogram (PCH) analysis, which can measure these fluctuations, assume an infinite Gaussian beam, which prevents their application to particles within domains bounded at the nanoscale. We therefore derived general forms of FCS and PCH for bounded systems. The finite domain form of FCS differs from the classical form in its boundary and initial conditions and requires development of a new Fourier space solution for fitting data. Our finite-domain FCS predicts simulated data accurately and reduces to a previous model for the special case of molecules confined by two boundaries under Gaussian beams. Our approach enables estimation of the concentration of diffusing fluorophores within a finite domain for the first time. The method opens the possibility of quantification of kinetics in several systems for which this has never been possible, including in the one-dimensional lipid tubules discussed in Part 2 of this paper.

**Statement of Significance:** Methods based on fluorescence measurements of molecular concentration fluctuations, including Fluorescence Correlation Spectroscopy and Photon Count Histogram analysis, are widely used to determine rates of diffusion, chemical reaction and sizes of molecular aggregates. Typically, the range over which the molecules can diffuse is large compared to the size of the focused laser beam that excites the fluorescence. This work extends these measurements to systems that are comparable in size to the excitation laser beam. This extends the application of these methods to very small samples such as the interior of bacterial cells or the diffusion of molecules along individual macromolecules such as DNA.

## 1. Introduction

Fluorescence correlation spectroscopy (FCS) and associated methods such as photon counting histogram analysis (PCH) have enabled the quantification of a range of kinetic processes, especially of processes related to diffusion (1). The basic experiment acquires the time course of fluorescence photons emitted by fluorophores that diffuse through a small focused laser beam that illuminates a tiny fraction of a much larger sample volume or area. The light emitted by random passage of the fluorophores across the laser-illuminated region fluctuates over time and can be analyzed statistically to estimate properties of both the diffusing molecules and the surrounding medium. The classical theories of PCH and FCS for interpreting these time courses are based on laser beams with Gaussian intensity profiles that extend over a quasi-infinite domain of the total sample volume even though most of the fluorescence is excited from a very narrow spatial range (1).

Although this is not a limitation for experiments on domains much larger than the beam waist, it precludes experiments in confined domains such as microfluidic channels whose sizes are comparable to the laser waist (2). In addition to such cases of confinement, there are important examples in which the illumination is not well modeled by a Gaussian, such as a single molecule experiment with a fluorescent DNA-binding protein diffusing along a DNA fragment that has a fluorescence quencher, donor or acceptor tethered to one end (3). To address this need for FCS and PCH analysis on systems with confined diffusion or non-Gaussian spatial illumination, we derived a general form of FCS and an associated PCH analysis for bounded systems under arbitrary illumination.

## 2. Background

### FCS

FCS analysis begins by gathering the time course of fluctuating fluorescence intensity, *I*(*t*), and computing the temporal autocorrelation function *G*(*τ*), which is the correlation of *I*(*t*) with itself when shifted by a time interval *τ*: 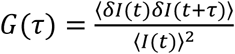, in which angle brackets indicate temporal averages, *t* is time, and the intensity fluctuation is *δI*(*t*) = *I*(*t*) − ⟨*I*(*t*)⟩. The intensity fluctuation relates to the concentration of fluorophores *c*(*r, t*) at position *r* and time *t*, the brightness per fluorophore, *B*, and the illumination profile *W*(*r*):

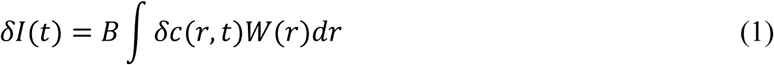

where *δc*(*r, t*) = *c*(*r, t*) − *C* is the local fluctuation of concentration around the macroscopic concentration *C.* Note that the product *BW*(*r*) is actually a combination of both illumination and detection sometimes called the detectable emission intensity distribution or spatial detectivity function. *B* and *W*(*r*) can be measured experimentally. For classic FCS, *W*(*r*) is approximated by a Gaussian, 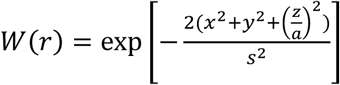, where *s* is the laser waist and *r_0_* is the position of the center of the laser beam and the autocorrelation function becomes 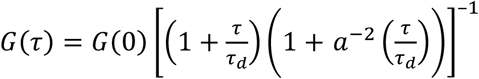 in which *a* is the ratio of the of axial to radial radii of the excitation spot, and the characteristic diffusion time *τ*_*d*_ is estimated by fitting this equation to experimental data.

This expression can be derived via a Fourier transform approach from the diffusion equation:

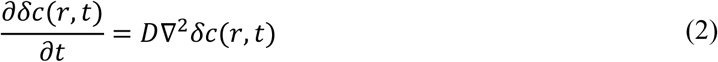

For an open system with no boundaries, the initial condition is that concentrations are uncorrelated spatially between any two points *r*′ and *r*, so that (1,4):

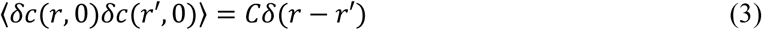

However, for bounded, finite systems this initial condition must be modified, and a series approach must be taken to solving the diffusion equation. We develop these approaches in the next section.

### PCH and FIDA

The analysis of photon counts/intensity was first discussed by Qian and Elson in early 90s(5, 6). It was further discussed by two papers in 1999: photon counting histogram analysis (PCH) (7), and fluorescence intensity distribution analysis (FIDA) (8). These two techniques are related mathematically (9), with the generating function in FIDA representing the Fourier transform of the PCH. Another important difference lies in the derivation procedures. PCH begins with calculation of the histogram for one molecule. Fluctuations for one molecule are due to the unevenness of the illumination and detection. The histogram for *n* molecules is the convolution of the histogram for one molecule *n* times. FIDA, on the other hand, begins with consideration of the fluctuations of fluorophores and photon counts from a small volume, *dV*. The overall distribution of the photon counts is the convolution of the distributions for all *dV*s, leading to the logarithm of the generating function of the photon counts and therefore an integration over *V*.

Below, we compare the two strategies for PCH and FIDA, which are convolutions of molecules and convolutions of space, respectively. PCH theory assumes that the distributions for different molecules are independent of each other, while FIDA theory assumes that the distributions for different positions are independent of each other. Although these two procedures give the same result for open systems, we show that FIDA is not suitable for bounded systems.

## 3. Methods

### Initial conditions for bounded FCS

The initial condition for FCS in an unbounded system is ⟨*δc*(*r*, 0)*δc*(*r*′, 0)⟩ = *Cδ*(*r* − *r*′), which states that concentrations (and hence numbers of molecules in a local volume) are initially uncorrelated. This further requires that when *r* = *r′*, the variance of *c* at *r*, equals its mean and therefore they follow a Poisson distribution. However, this initial condition does not apply in bounded systems because the limited number of molecules requires that concentration fluctuations at two different locations be negatively correlated; elevations of concentration at one location require reductions at other locations. As shown below, this leads to a binomial rather than Poisson distribution of molecules, thus a new initial condition.

The derivation begins by considering a 1-D bounded system with a length of *d* and *N*_*total*_ molecules, so that *C* = *N*_*total*_ /*d*. An arbitrary specific point *r* = *r*′, and all other points (*r* ≠ *r*′). In order to get ⟨*δc*(*r*, 0)*δc*(*r*′, 0)⟩, we will derive ⟨*δn*(*r*, 0)*δn*(*r*′, 0)⟩ for two cases: point *r* = *r*′ and *r* ≠ *r*′ (*n* is the number of molecules in a small interval Δ*r*).

First, for *r* = *r*′, the variance of concentration can be calculated as follows: for a small interval Δ*r*, each molecule has an equal probability of *p* = *Δr*/*d* that it can be in Δ*r* (therefore a probability of 1 − *p* that it does not reside in Δ*r*). Given a total number of *N*_*total*_, the number of molecules *n* in a small interval Δ*r* follows a binomial distribution in bounded systems and the variance is ⟨(*δn*)^2^⟩ = *p*(1 − *p*)*N*_*total*_, which is also the fluctuation of *n* at position *r* and time *t*=0:

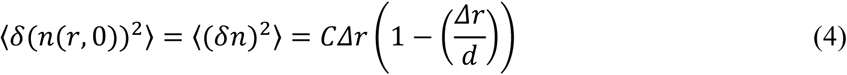

Second, for any other point (*r* ≠ *r*′), ⟨*δn*(*r*, 0)*δn*(*r′*, 0)⟩ needs to be derived. In order to do so, we considered the distribution of the sum of the number of molecules at *r* and *r*′, *n*(*r, r*′, 0) = *n*(*r*, 0) + *n*(*r*′, 0). It is worth noting that *n*(*r, r*′, 0) is not the 2D joint distribution but rather the distribution of the sum of two random variables which are supposed to be negatively correlated. Therefore, the variance of *n*(*r, r*′, 0) = *n*(*r*, 0) + *n*(*r*′, 0) is

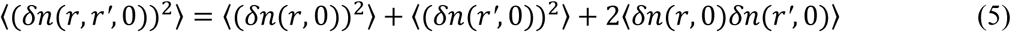

We are seeking the last term on the right side. Because ⟨*δ*(*n*(*r*, 0))^2^⟩ and ⟨*δ*(*n*(*r′*, 0))^2^⟩ have already been derived as in Equation (4), the only unknown term is the left side, ⟨*δn*(*r, r′*, 0)^2^⟩. To determine its expression, we consider the two small intervals at *r* and *r*′. As above, each molecule has a probability of *p* being at either *r* or *r*′ and a probability of 1 − *p* and being at somewhere else. Therefore, the molecules still follow a binomial distribution. Since the interval size increases two-fold (Δ*r* becomes 2Δ*r*), *p* becomes twice as big too: *p* = 2*Δr*/*d*. Therefore, its variance is:

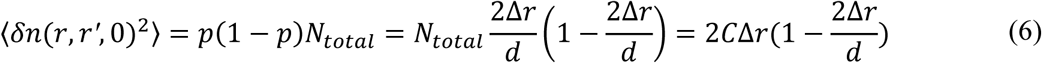

Rearranging Equation (5), substituting Equations (4) and (6):

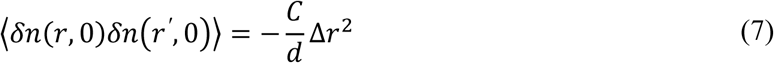

At last, rewriting Equations (4) and (7) in terms of the local concentration, *c* = *n*/*Δr, C* = *N*_*total*_/*d*:

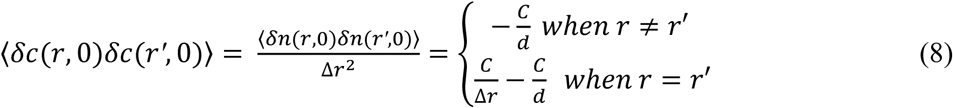

Given a function 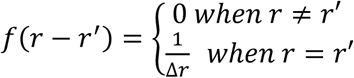 it can be shown that *f* is a delta function by proving the integral 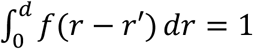 and 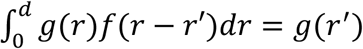. This can be easily done as Δ*r* and *dr* cancel out (These two integrals will be used in the derivation in the Appendix). Therefore ⟨*δc*(*r*, 0)*δc*(*r′*, 0)⟩ can be written in a simpler form, which is a delta function subtracted by a constant:

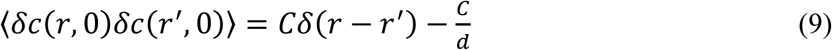

### Solution for bounded FCS

Solving the diffusion equation (Equation (2)) over a finite domain was achieved using a Fourier series approach. In this particular case, with Neumann boundary conditions, cosine series were used: 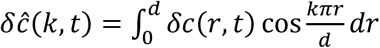, in which *δ ĉ* (*k, t*) is the Fourier transform of *δc* with wave number *k.* This leads to a solution of the diffusion equation:

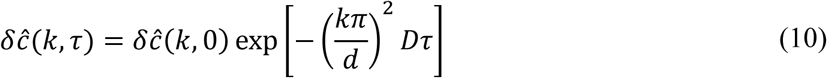

The unnormalized autocorrelation function can then be obtained (see Appendix for details):

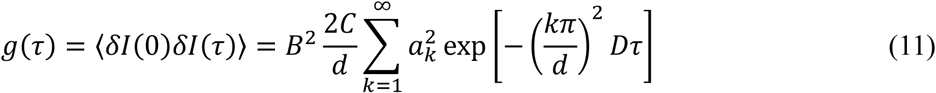

while

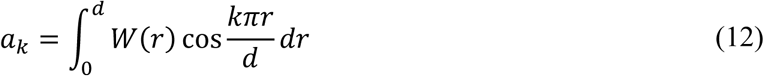

The normalization factor can be easily obtained as 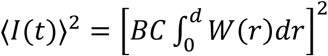. Therefore, the normalized form of the correlation function is

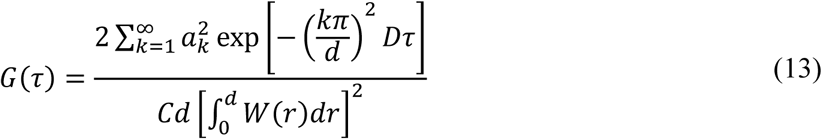

With the input *W*(*r*) and size *d*, this equation can be used to fit an experimentally determined autocorrelation function, as in classical FCS. The diffusion coefficient *D* and the concentration *C* can then be obtained.

It should be noted that when *W*(*r*) is constant, the autocorrelation *g*(*τ*) equals zero. This can be easily understood because under uniform illumination, there will be no fluctuations other than shot noise, which has no time correlation. For classic FCS in infinite systems, two sources of fluctuation other than shot noise exist: fluctuations caused by concentration fluctuations, and fluctuation caused by nonuniform illumination and detection. However, for the framework developed here for FCS in bounded systems, only the second source contributes.

### Bounded PCH and FIDA

As we have discussed earlier, PCH and FIDA have been developed to measure the number and brightness in fluorophore populations within an effectively unbounded region of solution. Although these two procedures give the same result for open systems, FIDA is not suitable for closed systems. The reason is that the distribution for different positions are not independent from each other for bounded systems. For open systems, the distribution of the number of particles at one position is a Poisson no matter how many particles there are in other positions. However, in the bounded system, the total number of particles is constant, and all the positions compete with each other for particles. Therefore, the convolution for all *dV*s is not suitable.

For one molecule in a bounded system, the photon counting distribution is,

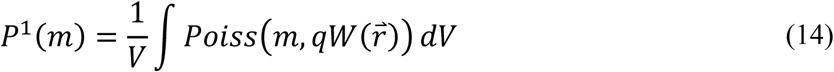

where *m* is the number of photons. For n particles, *P*^*n*^(*m*) is the self-convolution of *P*^1^(*m*) by *n* times (7). Using the generating function, the time-consuming convolution can be avoided. The generating function is

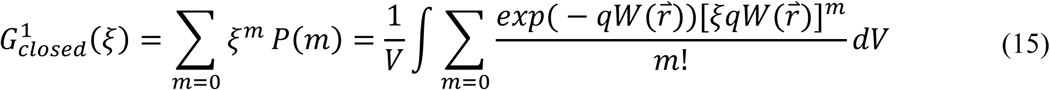

Finally, this yields

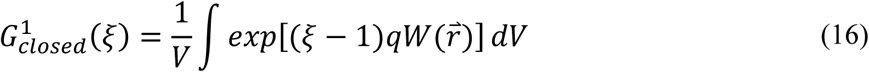

It can be easily shown that the distribution goes to a Poisson when 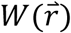 is constant over 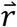, meaning the fluctuation is due only to the shot noise. For *n* molecules 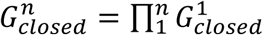. Given *ξ* = *e*^−*iω*^, inverse Fourier transform can be used to calculate the probability density function *P*(*m*).

## Results and Discussion

### Finite-Domain FCS reduces to previous results for special cases

To test our theory, we calculated *G*(*τ*) for the case of molecules diffusing in a 1D system, bouncing back and forth between two walls. Then, we compared our results to that given by Sanguigno’s method, which can be used only for Gaussian beams. As depicted in Figure 1, our calculation matches Sanguigno’s method very well for the special case of a Gaussian beam.

**Figure 1.**
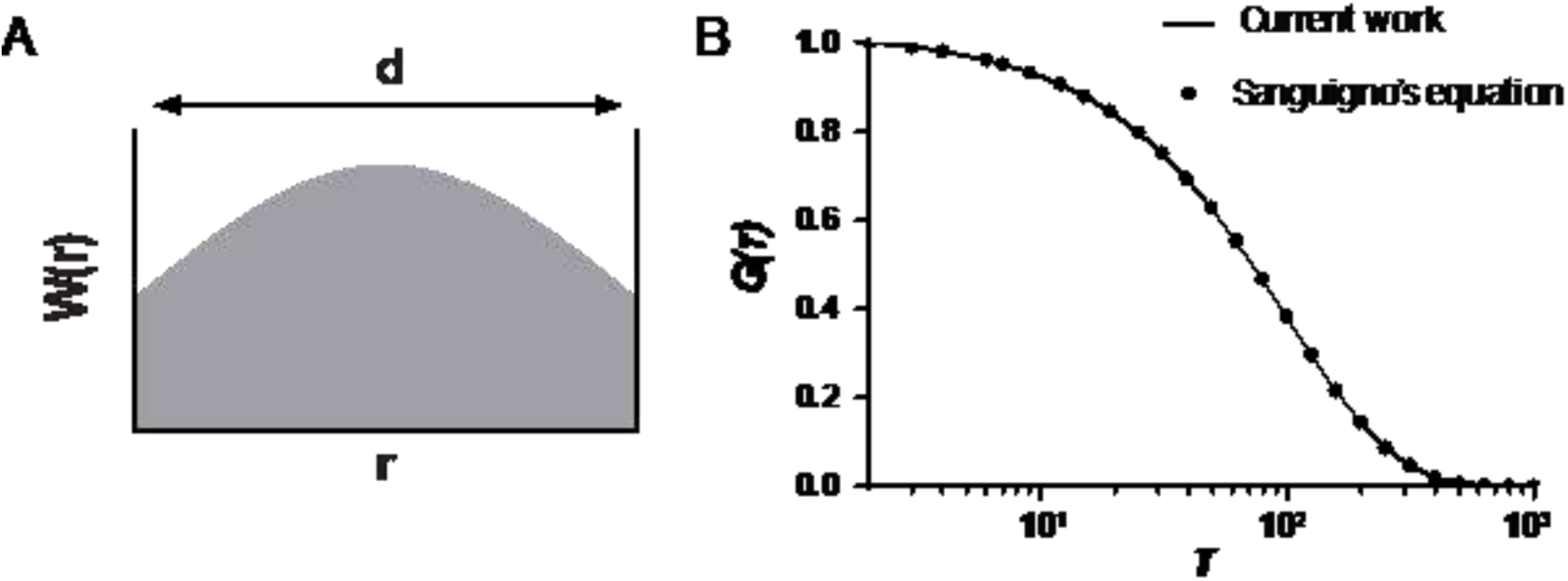
FCS in a bounded 1D system under a Gaussian beam focused at the center of the system. (A) Illustration of the system. System length d = 2, Gaussian beam waist is 1, diffusion coefficient D = 0.001, and the total number of particles is 1, which leaves the concertation C = 0.5 (all parameters are dimensionless). (B) Comparison with Sanguigno’s results. The correlation functions are divided by G(0). For Sanguino’s method, the G(∞) must be subtracted to force zero at the infinite time before dividing by G(0).

### Finite-Domain FCS matches simulation data for more general cases

Sanguigno’s calculation involves two extra steps: subtracting *G*(∞) and dividing by *G*(0) (We use the term “divide” instead of normalize to distinguish it from the normalization by the ⟨*I*(*t*)⟩^2^). The consequence of these extra steps is that the amplitude of the correlation is discarded. For traditional FCS, the amplitude has the information of particle concentration. High concentration leads to low correlation amplitude because the fluctuations will average out. However, there is no concentration term in Sanguigno’s derivation. When comparing the raw calculation (no subtraction and division) to simulated data, our result fit the data very well while Sanguigno’s result does not fit the simulated data (Figure 2A, dashed line and the inset)

**Figure 2.**
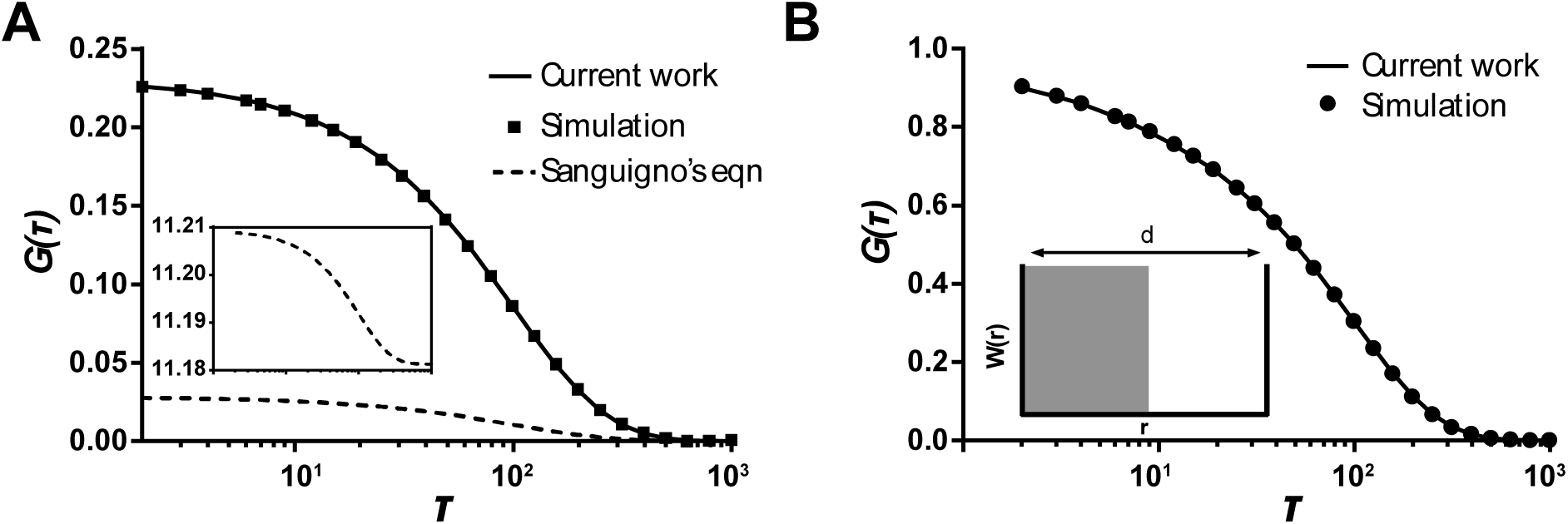
(A) Comparison between Sanguigno’s results and the current work fitting simulated data under a Gaussian beam. The inset panel shows the raw result by Sanguigno’s equations without the subtraction and division steps. The dashed line under the inset shows Sanguigno’s result that is only subtracted by G(∞) but not divided by G(0). The solid line shows our result without any extra steps. (B) Result for a square beam instead of a Gaussian beam, with which the left half of the system is evenly illuminated and the right half is not at all illuminated.

In Sanguigno’s method, the calculated correlation function does not approach zero at the infinite time. That value must therefore be subtracted to force it to be zero at the infinite lag time. When deriving our equations, we had the same problem until we found that the initial condition is not a simple Delta function anymore in bounded systems. Following the same derivation steps in the appendix, it can be easily proved that with the initial condition ⟨*δc*(*r*, 0)*δc*(*r′*, 0)⟩ = *Cδ*(*r* − *r′*), the *G*(∞) will be 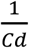 or 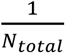 instead of zero, while *N*_*total*_ is the total number of particles in the system.

Compared to Sanguigno’s method, our method has two more advantages. First, it can be used for any other laser beam shape. Figure 2B shows that our theoretically calculated curve fits the simulated data very well for a square laser beam. Secondly, our method requires much less computational power. Using MATLAB, under the same conditions, our method improved the computation speed about 3000-fold. This is mainly because an error function of complex numbers in Sanguigno’s equations takes very long time to evaluate. Random walk simulation was also timed as a comparison.

**Table 1.** Time required to compute G(*τ*) using Sanguigno’s method, our method and direct simulation. For both Sanguigno’s and our method, the infinite sum was limited to only the first 100 terms (wavenumber) without loss of generality. The simulation consists of 10 million steps of random walk with Gaussian step-size.

|  | Sanguigno's method | Current work | Simulation |
| --- | --- | --- | --- |
| Computation Time | $96 \pm 5s$ | $0.026 \pm 0.006s$ | $3.6 \pm 0.1s$ |

### Finite Domain FIDA and PCH

We next assessed whether the approach could be used to determine the numbers and brightnesses of multiple populations of fluorophores diffusing through a laser beam in a confined space. In Figure 3, we calculated the PCH instead of correlation function using the same simulation described in Figure 1. We show that our calculations fit the simulated PCH very well. At the same time, results reveal that as the number of particles increases, the distribution approaches a Poisson distribution (Figure 3B, *N*=5, *B*=1).

**Figure 3.**
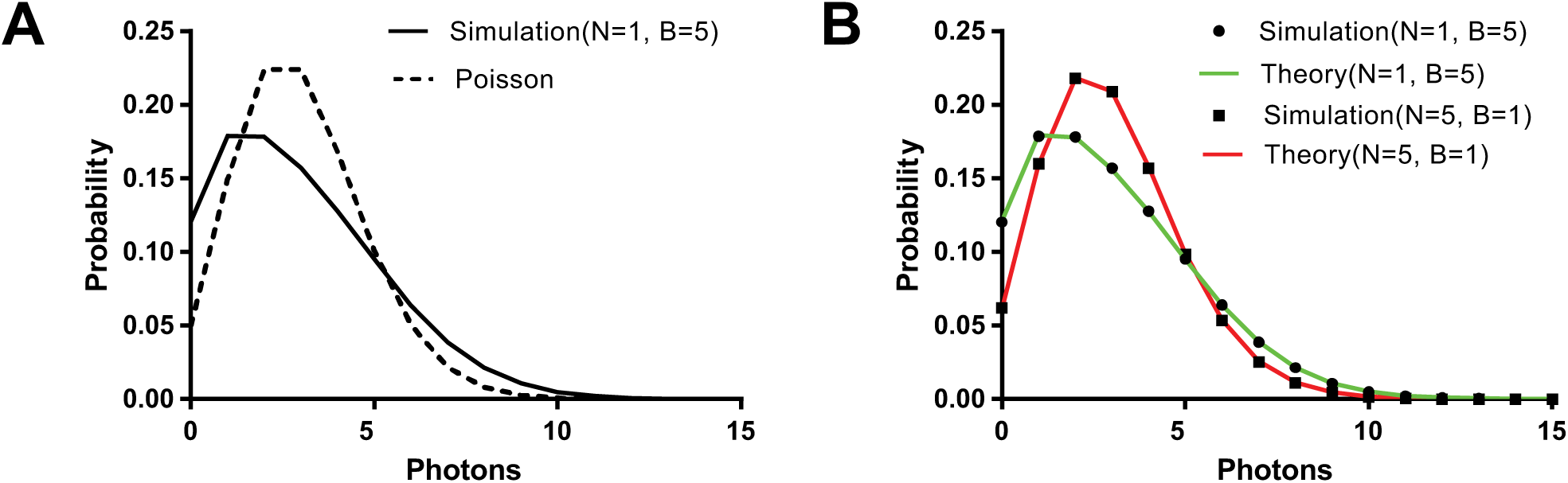
Photon counting histogram in bounded systems. (A) The histogram deviates from Poisson (with the same mean). (B) Compare the simulated data with the result calculated using the Fourier transform of Equation (16). Two cases were simulated while both give the same average intensity, which is the product of the number of particles N and their brightness B.

#### Conclusions and Perspectives

The theory for reflective boundaries can be useful for some single molecule studies, as depicted in Figure 3. Here, we give two potential applications. The diffusion of proteins along DNA is usually studied with very long DNA molecules whose length are much longer than the optical resolution. To obtain the diffusion rates, a fitting step is required to get the precise position of the proteins. This is also the basic principle of the STORM and PALM super-resolution microscopy. One of the drawbacks is that the fitting step is time consuming. Here, we propose a single molecule experiment in which short DNA can be used (Figure 4A). For this experiment, a pair of FRET fluorophores are conjugated on the proteins and the end of the DNA, respectively. The diffusion of protein leads to a change of the distance between the FRET pair and causes fluctuations of the detected intensity. The correlation function can be calculated and fitted with our theory for the reflective boundaries. In this case, the illumination function is 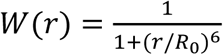, in which *R*_0_ is the Foster distance of the FRET pair. The DNA length can affect both the amplitude and the decay time of the correlation function (Figure 4A). Moreover, the correlation function is very different from the 1D FCS equation 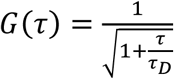 (Figure 4B).

**Figure 4.**
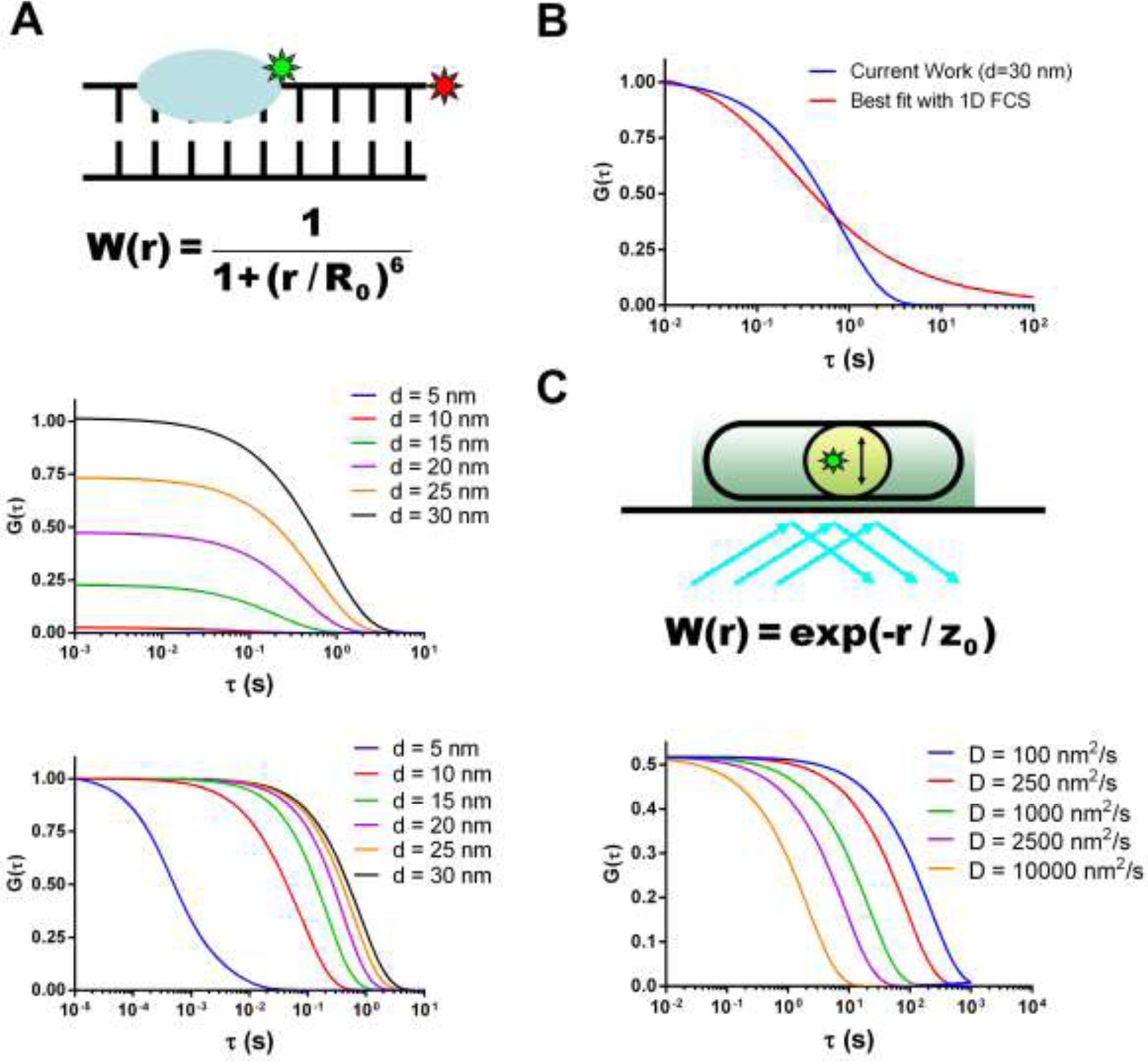
Examples of single molecule experiments for which our theory could be useful. (A) protein diffusion along short DNA molecules of different lengths. The middle and bottom subpanels show the curves without and with dividing by G(0), respectively. The diffusion coefficient D=100 nm^2^/s. (B)Results of fitting the 30 nm data in (A) with 1D FCS, 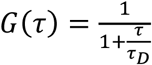. (C) Diffusion in a confined region in the cell studied using TIRF (total internal reflection fluorescence microscopy). Different diffusion rates were simulated. z_0_ is set to be 150 nm. The unit of D is nm^2^/s.

Bacterial cells provide another popular target for single molecule studies. A typical *E. coli* cell spans 2 µm and has a diameter of 1 µm. The diffusion of molecules in the cells is usually measured using TIRF (total internal reflection fluorescence microscope) in the x-y directions, which are parallel to the surface. However, if the diffusion is confined in a region that is smaller than the optical resolution, it is difficult to obtain the diffusion rates. Here, we proposed a method to study diffusion in the z-direction (in terms of microscope stage instead of cell geometry) taking advantage of the decaying excitation intensity for TIRF(10). As shown in Figure 4C, as the protein moves in the z-direction, the brightness will change. The intensity can be recorded, and the correlation function can be calculated and fitted to our theory. In this case, the illumination intensity decays exponentially, so that *W*(*r*) = *exp* (− *r*/*z*_0_).

To summarize, we derived the equations for FCS in bounded systems. These equations are applicable for any illumination function and could be potentially useful for many experiments. We also want to emphasize that the initial condition we found for solving the diffusion equations might be useful not only for FCS but also for other theoretical and experimental investigations.

## Appendix Derivation of the correlation function for confined FCS

The fluorescence intensity from a region illuminated by a laser is calculated as:

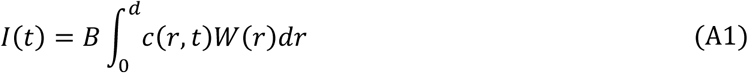

where *c*(*r, t*) is the concentration of fluorophores at position *r* and time *t, B* is the brightness per fluorophore, and *W*(*r*) is the illumination profile.

In response to a perturbation in concentration *δc*(*r, t*), the intensity fluctuates by an amount:

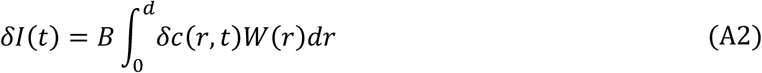

Therefore, the autocorrelation function can be written as

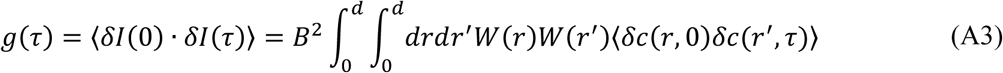

The autocorrelation function *g*(*τ*) can be computed from experimental observations because the intensity fluctuation can be measured. Because all terms in Equation (A3) can then be measured other than ⟨*δc*(*r*, 0)*δc*(*r*′, *τ*)⟩, it is possible to use Equation (A3) to calculate this. To relate these fluctuations to the diffusion coefficient, we solved the diffusion equation:

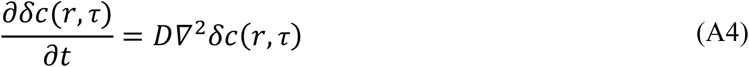

Our procedure for this involved a Fourier space approach. Solution of the diffusion equation in the Fourier space is:

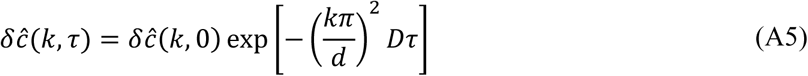

When *k* = 0, this becomes:

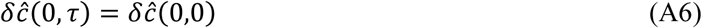

For a reflective boundary, the cosine Fourier transform of *δc* is:

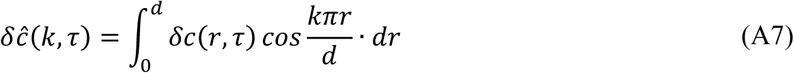

Note that at time *τ =* 0,

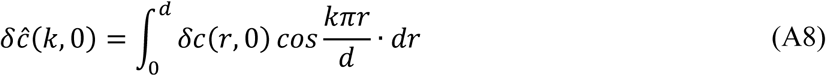

and, when *k* = 0,

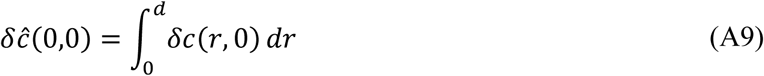

Substituting Equations (A6)-(A9) and taking the inverse Fourier transform to transform the solution into real space and obtain *δc*(*r*′, *τ*), we arrive at:

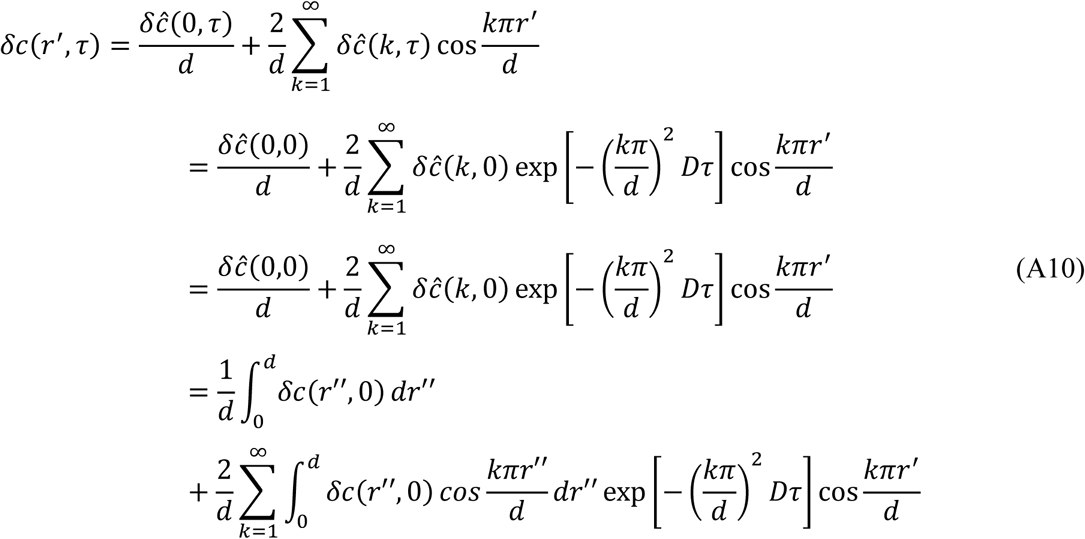

Hence,

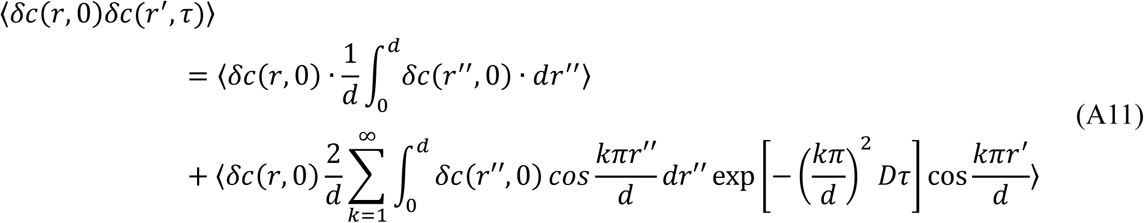

Rearranging by moving the angled brackets that represent time averages:

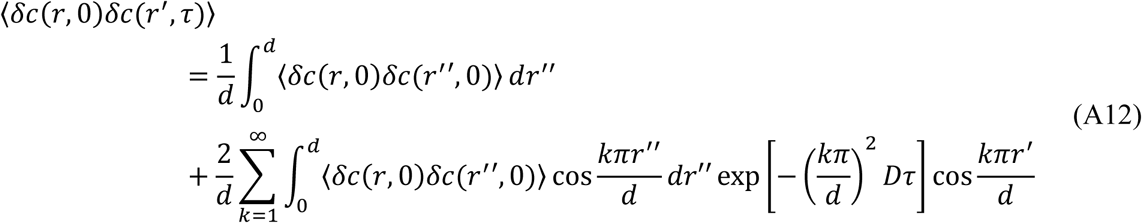

The next step is to substitute the initial condition 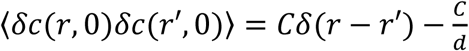 in equation (A12). To this end, the most two important properties of Delta function are used. They are: 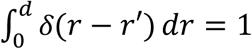 and 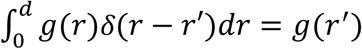.

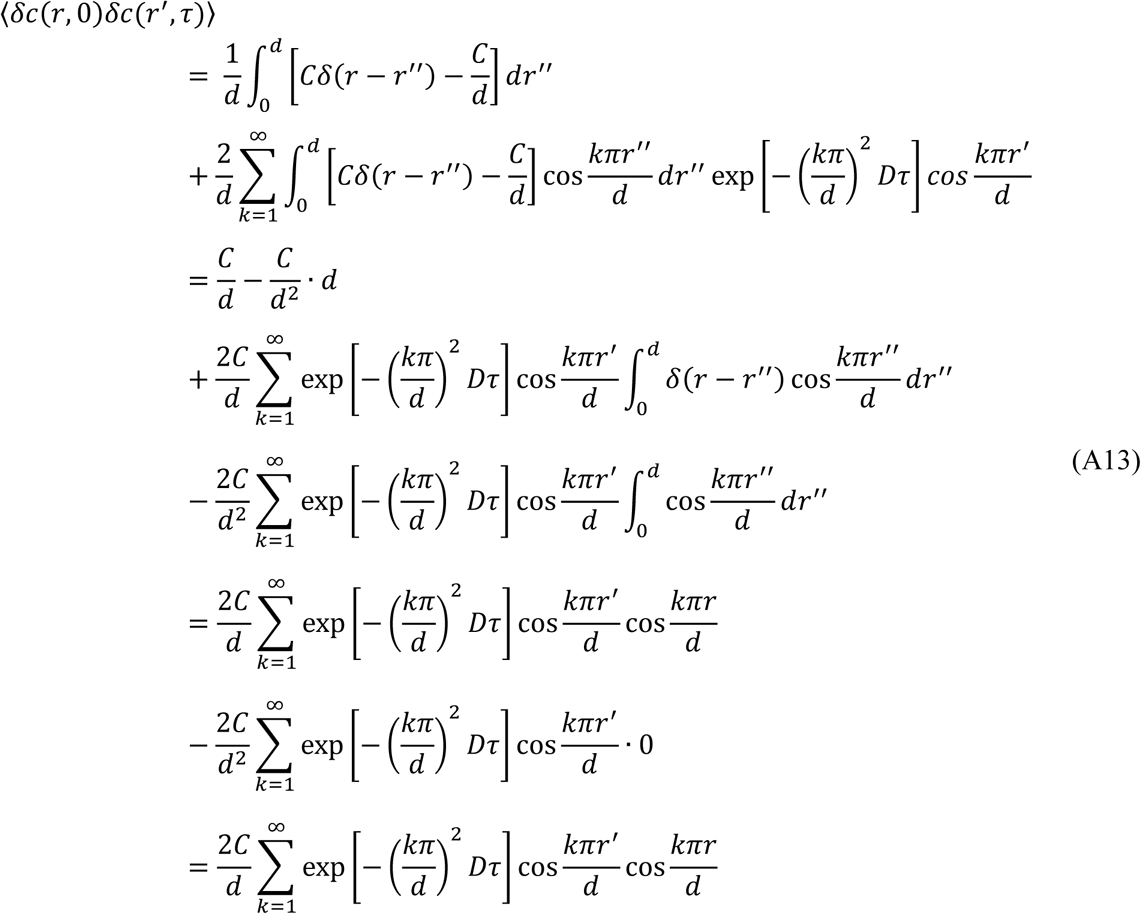

Substituting this expression for ⟨*δc*(*r*, 0)*δc*(*r*′, *τ*)⟩ into the autocorrelation function of Equation (A3) yields an expression that can be fit to estimate the diffusion coefficient *D* from experimental data:

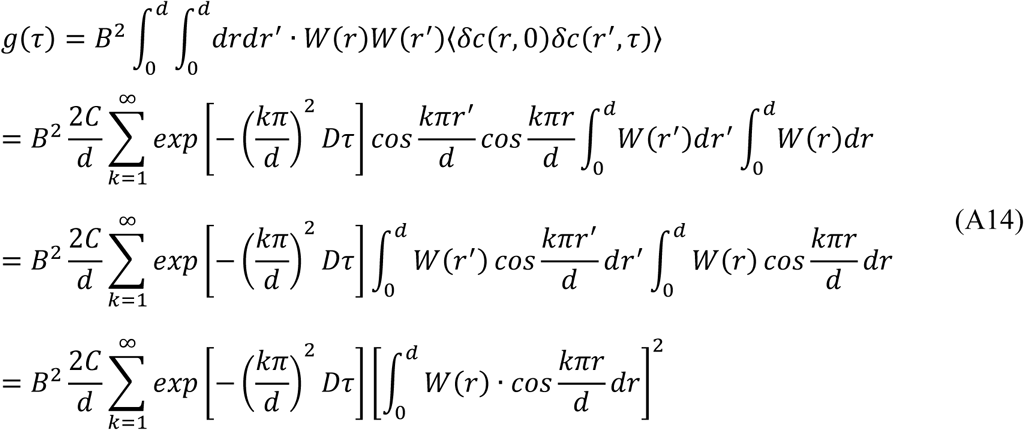

## Author Contributions

YJ conceived of and executed the study. AM, BX, ELE, GMG, provided counsel and assisted with the interpretation and presentation of the work. All authors contributed to writing the final paper..

## Acknowledgments

This work was funded in part by the NSF Science and Technology Center for Engineering Mechanobiology, grant CMMI 1548571. The authors thank Tony Pryse, Tim Lohman, and William McConnaughey for insightful commentary.

